# Project MinE: study design and pilot analyses of a large-scale whole genome sequencing study in amyotrophic lateral sclerosis

**DOI:** 10.1101/152553

**Authors:** Project MinE ALS Sequencing Consortium, Wouter Van Rheenen, Sara L. Pulit, Annelot M. Dekker, Ahmad Al Khleifat, William J. Brands, Alfredo Iacoangeli, Kevin P. Kenna, Maarten Kooyman, Russell L. McLaughlin, Bas Middelkoop, Matthieu Moisse, Raymond D. Schellevis, Aleksey Shatunov, William Sproviero, Gijs H.P. Tazelaar, Rick A.A. Van der Spek, Perry T.C. Van Doormaal, Kristel R. Van Eijk, Joke J.F.A. van Vugt, A. Nazli Basak, Jonathan D. Glass, Orla Hardiman, Winston Hide, John E Landers, Jesus S. Mora, Karen E. Morrison, Wim Robberecht, Stephen Newhouse, Christopher E. Shaw, Pamela J. Shaw, Philip Van Damme, Michael A. Van Es, Ammar Al-Chalabi, Leonard H. Van den Berg, Jan H. Veldink

## Abstract

The most recent genome-wide association study in amyotrophic lateral sclerosis (ALS) demonstrates a disproportionate contribution from low-frequency variants to genetic susceptibility of disease. We have therefore begun Project MinE, an international collaboration that seeks to analyse whole-genome sequence data of at least 15,000 ALS patients and 7,500 controls. Here, we report on the design of Project MinE and pilot analyses of newly whole-genome sequenced 1,264 ALS patients and 611 controls drawn from the Netherlands. As has become characteristic of sequencing studies, we find an abundance of rare genetic variation (minor allele frequency < 0.1 %), the vast majority of which is absent in public data sets. Principal component analysis reveals local geographical clustering of these variants within The Netherlands. We use the whole-genome sequence data to explore the implications of poor geographical matching of cases and controls in a sequence-based disease study and to investigate how ancestry-matched, externally sequenced controls can induce false positive associations. Also, we have publicly released genome-wide minor allele counts in cases and controls, as well as results from genic burden tests.

## Introduction

Amyotrophic lateral sclerosis (ALS) is a rapidly progressing, fatal neurodegenerative disease.^1^ Twin studies estimate the heritability of ALS to be ~60%, suggesting a strong genetic component contributing to disease risk,^2^ and approximately 10-15% of patients have clear family history disease.^3^ of Genetic risk factors have been extensively studied in familial ALS cases, and this effort has led to the identification of highly penetrant causal mutations, including mutations residing in *SOD1*,*^4^ TARBP*,*^5^ FUS*,^*6*^ and *C9orf72.^7,8^* In so-called sporadic ALS cases, who have no known family history of disease and comprise the majority of all cases, only a small number of other common genetic risk loci have been identified. Among these loci are the *ATXN2* CAG repeat expansion,^9^ mutations in *C21orf2*,^*10*^ common variation within the *UNC13A*,*^11^ SARM1*,*^12^ MOBP*,*^10^ and SCFD1^10^* loci and, most importantly, the highly pathogenic *C9orf72* repeat expansion. The latter highlights the typical categorization of “familial” and “sporadic” ALS as likely non-distinct groups with overlapping genetic architectures.

Despite these recent advances in ALS genetics, the bulk of risk loci in ALS remain undiscovered. The most recent and largest ALS GWAS, performed in 12,577 cases and 23,475 controls, showed a disproportionate contribution of low-frequency variants to the overall risk of ALS.^10^ Genome-wide association studies to date have focused almost exclusively on common variation (minor allele frequency > 1%), leaving lower-frequency and rarer variants segregating in the general population essentially untested. Thus, there is a pressing need to study rare variation across the full length of the genome in cases with and without family history of disease. To this end, we have begun Project MinE, a large-scale whole-genome sequencing study in ALS. The project leverages international collaboration and recent developments in sequencing technologies, allowing us to explore the full spectrum of genetic variation in samples collected worldwide. Project MinE seeks to obtain sequencing data in 15,000 ALS patients and 7,500 matched controls with the aim of identifying new loci associated to ALS risk, fine-map known and novel loci, and provide the ALS community (and the disease research community at large) with a publicly-available summary-level dataset that will enable further genetic research of this and other diseases. A data access committee controls access to raw data, ensuring a FAIR data setup (http://www.datafairport.org).

While common variant association studies have become mostly standardized over the last decade, rare variant association studies such as Project MinE face an array of new challenges. Sequencing studies demand large sample sizes to detect small effects at rare variants, thus making large-scale collaborative consortia a necessity. Sequence data itself, measuring into the terabytes, poses a substantive data storage, processing, and management challenge. The analytic effects of (tight) spatial clustering of rare variants is not yet well understood,^13,14^ making careful case/control selection and proper handling of population stratification key. Rare variant association studies typically employ genic burden testing to overcome power problems, thus requiring a series of analytic choices be made regarding the functional annotation^15^ and statistical analysis of the data. Here, we discuss the “pilot phase” of Project MinE, performed in 1,264 cases and 611 controls collected in the Netherlands. Though the sample size is too small to detect moderate-effect genetic associations, it allows us to explore the implications of these many analytic challenges faced by Project MinE or any disease study employing sequence data, to understand the genetic basis of disease. We outline the formation of the consortium, study design challenges, data quality control and analysis approaches employed by the project, and publicly release all minor allele counts in cases and controls together with results from genic burden tests derived from this dataset.

## Material and methods

### Consortium design

The MinE consortium includes ALS research groups from 16 different countries collaborating in a “franchise” design (www.projectmine.com, **Figure 1**). This design means that samples and sample-affiliated data (e.g., dense phenotyping) from partners are collected, processed and stored according to the same protocol, while partners maintain full control over their samples, affiliated data, and any additional data generated on those samples. Within the consortium, ALS patients and controls are ascertained through clinics affiliated to the research groups. At these clinics, neurologists obtain a standardized core clinical data set with phenotypic information (**Supplementary Table 1**) and blood is drawn for DNA isolation. DNA samples are stored on site, but are collectively prepared for sequencing at commercial sequencing providers, in batches of up to 1,000 samples. The consortium continues to expand, welcoming research groups who wish to dedicate time and raise money to understand the genetic and environmental underpinnings of ALS.

**Figure 1.**
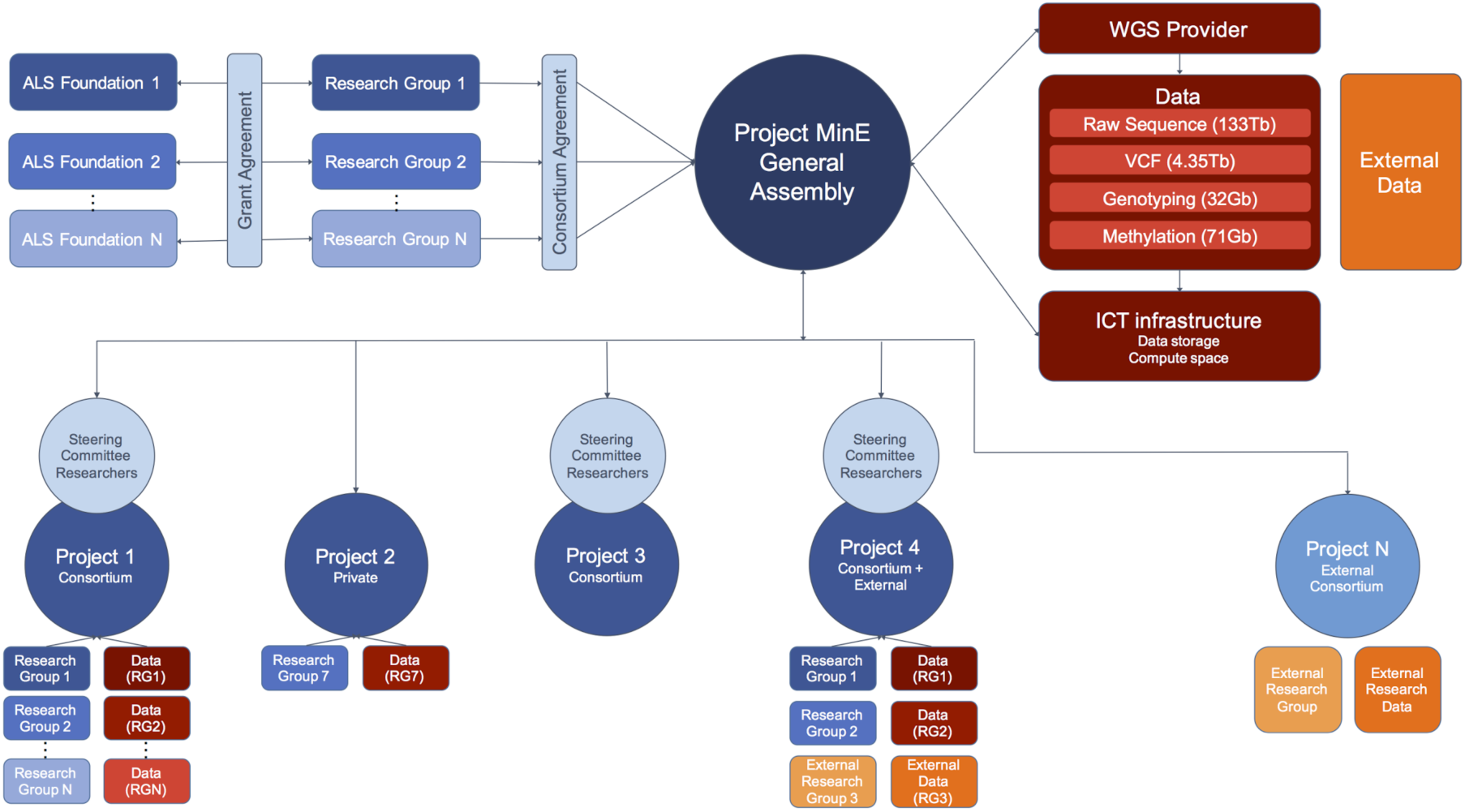
Consortium design. Consortium structure including funding agencies, research groups, the Project MinE general assembly, sub-projects, and data management. Each research group obtains funding using uniform grant proposals shared between the research groups. DNA samples are provided to the sequencing providers via the Project MinE general assembly so research groups can profit from a lower pricing scheme available when large numbers of DNA samples are provided together. Samples are sequenced, and data (wholegenome sequence data in raw and genotype-called format, SNP array data, and methylation array data) is centrally stored (the size of the data for the first 1,935 samples is indicated in parentheses; the full dataset for N = 22,500 samples will be ~1.7 petabytes). External data can also be contributed to and integrated into the dataset. Different research groups, including external groups, collaborate to work on specific projects. After review of the project proposal to the general assembly, researchers working on these projects will be granted access to the data and may also use the central ICT facilities. ALS: amyotrophic lateral sclerosis, WGS: whole genome sequencing.

A number of variables, including pricing for varying numbers of genomes, DNA input requirements, availability of PCR-free library preparation, and methods for data delivery, were all considered during selection of a sequencing provider (**Supplementary Table 2**). The majority of the data to date have been sequenced at Illumina (San Diego, CA, USA); a small subset is currently being sequenced locally in Australia and Canada. From Illumina, the data are transferred through the internet (via a secure connection) to SURFsara (Amsterdam, The Netherlands). After arrival, the data is automatically checked for corruption and stored in duplicate on two geographically separated tape silos. The storage system is connected to a distributed parallel file system (dCache^16^) that provides high performing file service to a 5,600 core high-throughput computing cluster.

From here, consortium members can access the genomes of their samples and ask that their data be additionally backed up on the High Performance Compute cluster at the University Medical Center Utrecht (Utrecht, The Netherlands). Analysis teams (possibly including researchers from external groups) can submit analysis proposals to the consortium and then gain access to all genomes and make use of the SURFsara compute facilities (**Figure 1**). Access to the data or usage of the compute facilities is contingent on (a) X.509 authentication certificates and (b) proposed analyses for the data. This data infrastructure allows, for example, particular cohorts and individuals access to particular subsets or all of the data, depending on the analysis. Based on permission given by the manager of the data (i.e., the principal investigator of the contributing research group), access is provided through the gridftp protocol v2 that is tailored for large and reliable file transfers. The X.509 certificates allow for the possibility to scale out to other compute facilities; given valid credentials, the data can then be accessed from anywhere.

### Sample selection

Patients were diagnosed with definite, probable, and probable lab-supported ALS according to the revised El Escorial Criteria.^17^ For the pilot analyses presented here, all patients were seen by neurologists specialized in motor neuron diseases at the University Medical Center Utrecht and Academic Medical Center (Amsterdam, The Netherlands). The samples were all included in the Prospective ALS study in the Netherlands (PAN), an incidence-based registry of ALS cases^18^. Controls were population-based controls that were matched for age, sex, and geographical region. As is true for all contributing cohorts to Project MinE, cases and controls were ascertained in a roughly 2:1 ratio; the 2:1 case-control ratio was selected to improve detection of variants in cases and with the plan to include publicly-available sequenced controls in future analyses.

### Phenotypic information

For all participants in Project MinE, we have defined a core clinical dataset, meant to be collected and made available for all cases (**Supplementary Table 1**). Phenotypic information is stored physically separately (<u>https://euromotor.umcutrecht.nl)</u> from the genetic data in a clinical database called Progeny. The phenotypic and genetic data can be connected through sample identifiers. Phenotypic data storage is organized similarly to the sequencing data: every contributing group has default full access to their own phenotypic data, but data can easily be jointly shared analysed if desired. This core clinical dataset (**Supplementary Table 1**) includes date of birth, sex, site of onset (spinal or bulbar), date of disease onset, diagnostic category (according to both revised and original El Escorial scoring), Forced Vital Capacity at time of diagnosis, cognitive status measured by the Edinburgh Cognitive and Behavioural ALS Screen (ECAS), and a revised ALS functional rating scale score (ALSFRS-R).^19^ The required endpoint data include date of death, date of starting invasive ventilation or requiring continuous non-invasive ventilation. For the pilot data used in this study, we obtained additional information on place of birth through the Dutch Municipal Personal Records Database.

### Whole-genome sequencing

Venous blood was drawn from patients and controls from which genomic DNA was isolated using standard methods. We set the DNA concentrations at 100ng/ul as measured by a fluorometer with the PicoGreen^®^ dsDNA quantitation assay. DNA integrity was assessed using gel electrophoresis. All samples were sequenced using Illumina’s FastTrack services (San Diego, CA, USA) on the Illumina HiSeq 2000 platform. Sequencing was 100bp paired-end performed using PCR-free library preparation, and yielded ~40x coverage across each sample. The Isaac pipeline 20 was used for alignment to the hg19 reference genome as well as to call single nucleotide variants (SNVs), insertions and deletions (indels), and larger structural variants (SVs). Both the aligned and unaligned reads were delivered in binary sequence alignment/map format (BAM) together with variant call format (gVCF) files containing the SNVs, indels and SVs.^20^ gVCF files were generated per individual and variants that failed the Isaac-based quality filter were set to missing on an individual basis.

### Data analysis

#### Quality control

Quality control (QC) of the data included QC at an individuals and variant level. Full details of QC are provided in the supplement.

#### Principal component analysis

Principal components were calculated for all individuals including variants at different allele frequency thresholds using GCTA.^21^ The eigenvectors of the first twenty principal components were regressed on latitude and longitude of birthplace. In a leave-one-out scheme, this linear model was used to predict the birthplace for the individual left out.

#### Identity-by-descent analysis

We phased all non-singleton variants using SHAPEIT2^22^ and then used BEAGLE4^23^ to detect runs of identity-by-descent (IBD) between individuals.

#### Association analysis

Genotypes at common variants (MAF > 0.5%) were tested for an association with case/control status using logistic regression (PLINK v1.9)^24,25^ assuming an additive model. Optionally, the first 10 principal components (PCs) were included as covariates. To test rare variation, we performed genic “burden” testing. All variants were functionally annotated using ANNOVAR^26^. We then determined three functional groups for gene-based association testing: (a) loss of function (LOF) mutations (premature stop mutations, stop-loss mutations, mutations at splice sites, and frameshift indels), (b) nonsynonymous mutations, and (c) LOF and nonsynonymous mutations (aggregated). Burden testing (T1, T5, Variable Threshold, Madsen-Browning and SKAT) was implemented using ScoreSeq^27^ and performed across all variants with MAF < 5%. All burden tests were adjusted for sex and the top ten PCs and performed on the QC-passing set of unrelated samples (1,169 cases and 608 controls).

#### Population stratification and externally sequenced controls

To assess the type I error in a burden testing framework, we simulated 100 different phenotypes for two scenarios: perfect matching and imperfect matching with a North-to-South gradient for the number of cases (keeping the case/control counts the same, **Supplementary Figure 1**), and then performed burden testing on LOF and nonsynonymous mutations with and without principal components as covariates. The most extreme p-value from each simulation was extracted (resulting in 100 extreme p-values total, per scenario); the fifth-most extreme p-value represented the p-value threshold necessary to maintain study-wise type I error at 5%. Subsequently, we assessed the impact of including ancestry-matched, externally sequenced controls (the Genome of the Netherlands, GoNL^14^) instead of controls sequenced as a part of Project MinE. As their data is mostly available formatted as VCF files, we combined both datasets on a VCF level, without realignment or joint variant calling. The structure of this dataset was described by principal component analysis and we assessed inflation of the test statistics when externally sequenced controls were included in genome-wide burden testing.

### Study approval and informed consent

All participants gave written informed consent and the institutional review board of the University Medical Center Utrecht approved this study. Additional approval was obtained to access the Dutch Municipal Personal Records Database.

## Results

### Baseline characteristics and description of data

After quality control, 1,169 unrelated Dutch-ancestry cases and 608 ancestrally-matched controls were available for analysis. Their baseline characteristics are displayed in **Table 1** and the distribution of cases and controls throughout The Netherlands is displayed in **Figure 2a**. In total, 42,200,214 SNVs and indels passed quality control. The majority (69%) of these sequenced variants were rare (MAF < 0.001, < 3 allele observations in our dataset, **Supplementary Figure 2**). In particular, the bulk of these rare variants were not observed in publicly available datasets of whole-genome sequencing data including the Genome of The Netherlands project (GoNL release 5 from 28-10-2013, N = 498) and the 1000 Genomes Project (Phase 1 from 23-11-2010, N = 1,093). This observation reflects population-specific variants and the growing number of rare variants that will continue to be discovered as sequencing is performed in increasingly larger samples around the globe. As expected, most common variants (MAF > 1%) have been observed in these two datasets (97.5% in GoNL and 98% in the 1000 Genomes Project) reflecting global sharing of common variation.

**Figure 2.**
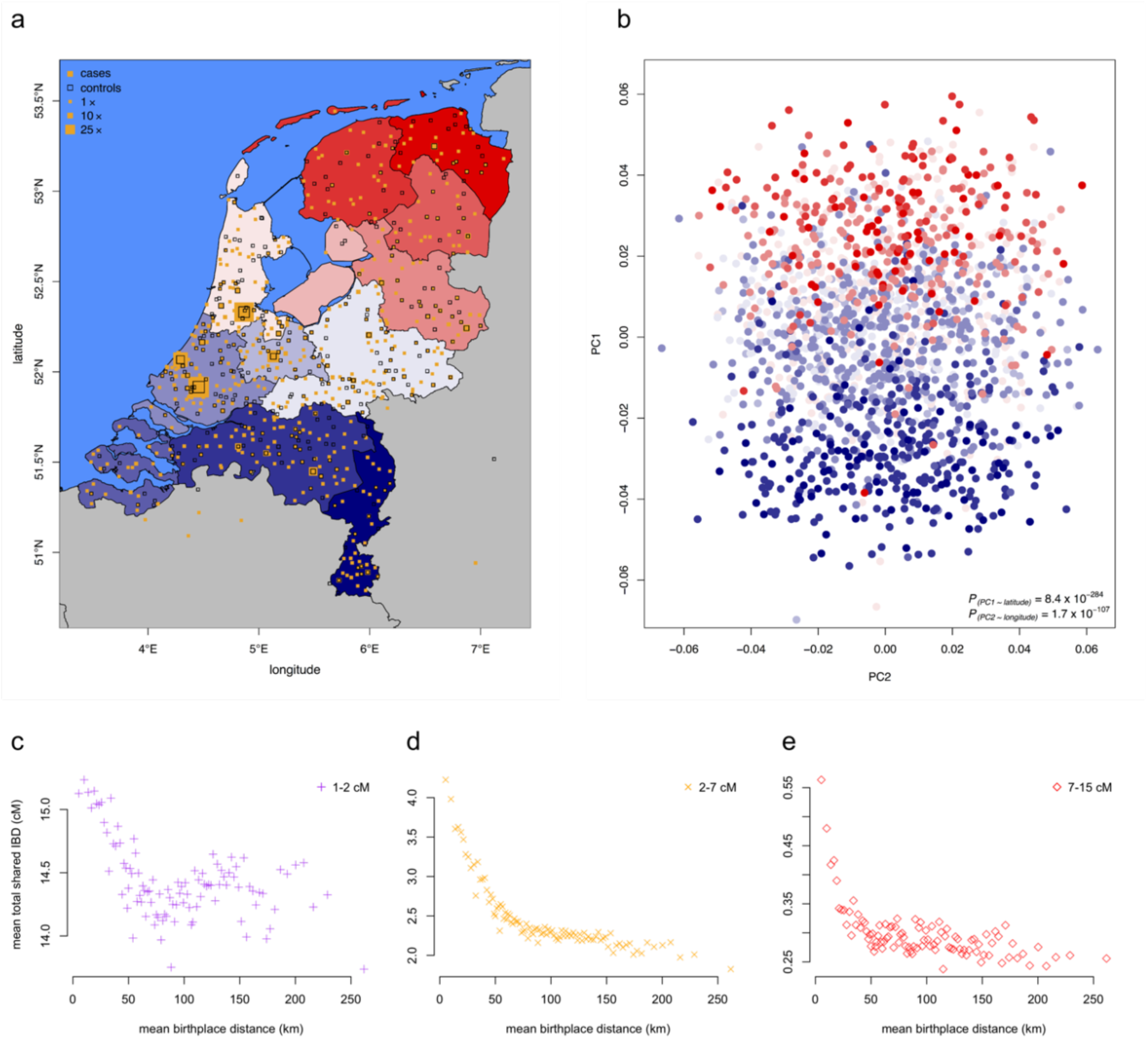
Population structure by principal component analysis. **(a)** Birthplaces of cases and controls for individuals born in The Netherlands. **(b)** The first two principal components reflect the geographical distribution of samples. Individuals are shaded by province of birth as shown in panel (a). The first principal component is strongly correlated with birthplace latitude (p = 8.4×10^-284^) and the second principal component with longitude (p = 1.7×10^-107^). **(c-e)** Relation between birthplace distance between pairs of individuals and shared IBD segments 1-2 cM **(c)**, 2-7 cM **(d)** and 7-15 cM **(e)**. Long IBD segments are, as expected, mostly shared by individuals born close to one another. PC: principal component, MAF: minor allele frequency, km: kilometer, cM: centimorgan.

**Table 1.** Baseline characteristics of samples included in Project MinE. WT = wild-type, IQR = interquartile range.

|  | ALS | Controls |
| --- | --- | --- |
| N | 1196 | 610 |
| sex, % female | 41.1 | 43.1 |
| Type |  |  |
| familial (%) | 31 (2.6) | - |
| sporadic (%) | 1165 (97.4) | - |
| *C9orf72* |  |  |
| WT (%) | 1095 (91.6) | - |
| expanded (%) | 74 (6.2) | - |
| unknown (%) | 27 (2.3) | - |
| age at onset, years (IQR) | 63.97 (56.58 - 70.57) | - |
| site of onset |  |  |
| bulbar (%) | 386 (32.3) | - |
| spinal (%) | 718 (60.0) | - |
| thoracic (%) | 30 (2.5) | - |
| unknown (%) | 62 (5.2) | - |
| survival, years (IQR) | 2.29 (1.58 - 3.28) | - |
| deceased (%) | 966 (80.8) | - |

### Geographic clustering

The first and second principal components reflected the geographical distribution of cases and controls in detail (**Figure 2b**). The eigenvectors of the first principal component explained 55% of the variance in latitude (p = 8.4 × 10^-284^) and the second principal component explained 24% of the variance in longitude (p = 1.7 × 10^-107^) of the geographical distribution of the samples across the Netherlands. Prediction models including the first twenty principal components predicted the birthplace of individuals with high accuracy (**Supplementary Figure 3**). The accuracy increased when including rarer variants; the median distance between the predicted and actual birthplace decreased from 50 kilometres (31 miles) when considering common variants (MAF > 0.1), to 36 kilometres (22 miles) when including rarer variants (MAF > 0.001), illustrating the strong geographical clustering of rare variants.^28^

Sharing of IBD segments showed strong geographical patterns throughout the Netherlands. As was observed in a previous population genetics study in The Netherlands there is a clear North-to-South gradient for sharing of shorter (and thus older) IBD segments (**Supplementary Figure 4a**). We also confirmed the finding that people born in the Southern provinces shared more of these IBD segments with people from the Northern provinces than they share with people within their own province, reflecting the previously described complex migration patterns^14^. Sharing of longer IBD segments, reflecting more recently shared ancestry, was highly dependent on the distance of birthplaces between individuals (**Figure 1c-e** and **Supplementary Figure 4b**).

### Association testing in ALS cases and controls

We found no variants reaching genome-wide significance (p = 5.0 × 10^-8^) with ALS in our sample, as expected given the sample size. We looked up known associated single-variants, three of which showed nominal association to the trait (p < 0.05, **Supplementary Table 1**). Similarly, no loci achieved exome-wide significance in burden testing (p ≈ 5.0 × 10^-7^, after adjusting for 20,000 tested genes, the various burden test types, and sets of SNPs tested; **Supplementary Figure 5**) due to limited power (**Supplementary Figure 6**). Burden testing p-values for all genes tested can be found at http://databrowser.projectmine.com/.

### The implications of study design

We sought to evaluate how matching cases and control, thereby introducing fine-grained population structure, influenced the burden testing type I error rate. Therefore, we simulated phenotypes that (1) created perfectly geographically matched case-control sets and (2) introduced population structure between patients and controls (**Supplementary Figure 1**). We subsequently ran simulations that did not correct for any covariates or those that included common-variant (MAF > 0.1%) principal components. Imperfect matching did not yield a markedly more stringent p-value to maintain a study-wide type I error at 0.05 for all burden tests (**Table 2**), likely due to limited power. Correcting for PCs did not affect the p-value to maintain study-wise type I error at 0.05 either.

**Table 2.** Populations structure and use of principal components (PCs) to control type I error. No marked differences were observed between the p-values in perfect or imperfect matching scenarios (with or without PCs) to maintain the study-wise type I error rate at 0.05. Common PCs = principal components calculated on all common SNPs (MAF > 0.01). T1, burden test that considers variants with MAF < 0.01; MB, Madsen-Browning burden test that inversely weights variants by frequency; VT, Variable Threshold test; SKAT, Sequence Kernel Association Test.

**Scenario 1: perfect matching**
|  | *Type I error in burden tests* |  |  |  |
| --- | --- | --- | --- | --- |
|  | *T1* | *MB* | *VT* | *SKAT* |
| No PCs | $4.25 \times 10^{-6}$ | $3.38 \times 10^{-6}$ | $3.94 \times 10^{-6}$ | $2.35 \times 10^{-6}$ |
| Common PCs | $2.86 \times 10^{-6}$ | $3.99 \times 10^{-6}$ | $8.12 \times 10^{-6}$ | $2.24 \times 10^{-6}$ |

**Scenario 2: imperfect matching**
|  | *Type I error in burden tests* |  |  |  |
| --- | --- | --- | --- | --- |
|  | *T1* | *MB* | *VT* | *SKAT* |
| No PCs | $4.40 \times 10^{-6}$ | $1.90 \times 10^{-5}$ | $1.46 \times 10^{-5}$ | $6.72 \times 10^{-6}$ |
| Common PCs | $5.35 \times 10^{-6}$ | $9.35 \times 10^{-6}$ | $1.04 \times 10^{-9}$ | $7.81 \times 10^{-6}$ |

To test the analytic implications of a study design that uses its resources solely for the sequencing of cases and collects controls from an external source, we merged genotypes from the ALS cases with Dutch-ancestry samples whole-genome sequenced as part of the Genome of the Netherlands (GoNL) project. ^14^ Principal component analysis indicated a clear separation of the two projects, where PC1 effectively captured each project (**Supplementary Figure 7a**). Removal of highly differentiated SNPs with a frequency difference > 5% across the two projects somewhat mitigated this separation (**Supplementary Figure 7b**). As expected, single-variant testing between ALS cases and GoNL controls revealed excessive genomic inflation (⍰ = 1.12). Similarly, burden testing comparing ALS cases and GoNL controls revealed a strongly inflated QQ plot and genomic inflation factor (⍰ = 2.49), demonstrating the challenges of using externally sequenced controls to identify disease genes in a separately-sequenced set of cases and controls.

## Discussion and future perspectives

Family-based and population-level genetic studies have revealed ALS as a complex disease with a distinct role for rarer genetic variation. Although numerous genetic variants have been identified as conferring ALS risk, the genetic basis in the vast majority of cases is not yet understood. Here, we have described the design and pilot analyses of a large-scale whole-genome sequencing study aimed at discovering new genetic risk factors and further elucidating the genetic basis of ALS.

### Data sharing and open-access science

Data sharing and transparency in scientific research is advantageous for a host of reasons: it allows for analyses across large sets of samples, particularly for lower-prevalence diseases such as ALS; ensures rigorous experiments that can be reproduced by external groups; and allows for publicly-funded research to be made available to the public itself. Project MinE cannot publicly release individual-level genotype data due to consent. However, genotype frequency information and genic burden testing results are publicly available at the project’s online browser (<u>http://databrowser.projectmine.com/</u>). This browser will be continuously updated as the project expands, and will provide more detailed data integration in future releases. To further enhance the transparency within the consortium itself and with external collaborators and researchers, the project has begun a GitHub repository, to share and maintain scripts and data processing pipelines.

### The advantages of ‘franchising’

Given the relatively low prevalence of ALS, international collaboration is crucial to assembling large samples. Thus far, the “franchise” design of our consortium has allowed us to rapidly expand to sixteen participating research groups globally. This joint effort has collected enough resources to sequence >7,000 samples, with more currently being collected. Ongoing analyses include case-control association testing, as well as association testing in age-of-onset, and survival time. Additionally, the available core clinical dataset, uniformly collected for all individuals included in Project MinE, will allow for a host of other discovery analyses.

### Study design implications: data management, population architecture, and external controls

The pilot analyses, though small in sample size, yield several immediate and important conclusions. First, while genotype-called data (in VCF format) is easily handled by a high-performance compute cluster, whole-genome BAM files (approximately 80 gigabytes per genome) demand a compute infrastructure that can handle terabytes and even petabytes of data. This includes direct delivery of the genomes that are currently being sequenced via a direct high-speed connection between the sequencing provider and a computer cluster with a GRID architecture (SURFsara). Though some research institutions are already prepared for such a deluge of data, others will need to carefully consider compute infrastructure and the technical ramifications of assembling big data before embarking on large-scale sequencing analyses. Further, having all data stored in a single location enables analysis across the full dataset, including on BAM-level data.

Second, initial quality control of the data suggests that the high-depth sequencing and downstream genotype calling is yielding high-quality data, reflected in our ability to reconstruct, through principal component analysis and identity-by-descent analyses, demographic observations that have been made in separate large-scale sequencing efforts in the Dutch population. ^14^ Further, the population analyses performed here and in particular the birthplace analysis, highlights the sensitivity of genome-wide sequence data. The data must be carefully protected such that those who have donated DNA to enable disease research, will also remain anonymous (in accordance with the consent provided).^29^

Third, the high-resolution geographical clustering of rare variants as observed in these analyses could pose challenges in finding rare variants that contribute to ALS risk. The commonly applied burden test approach, however, does not seem to suffer from increased type I error rates induced by modest imperfections in geographically matching samples, as was shown by the burden test simulations. While it is possible that aggregating rare variants (found in potentially many geographic locations) in these tests makes the approach more robust to population structure, these results should be interpreted with caution. The modest power in our pilot sample likely limited our ability to detect the increasing type I error rates induced by imperfect geographic sample matching. With the increasing number of ancestrally diverse samples that are being included in Project MinE, false positive associations due to population stratification remain a serious concern, especially when our aim becomes identifying the causal single variant(s) that drive the genic burden signal. Correcting for principal components can help to control type I error, though potentially not sufficiently so in burden testing. More sophisticated analytic approaches, such as employing a linear mixed model in single variant and genic burden testing, will likely be helpful in controlling for population stratification across the full set of Project MinE samples.^30^

We further demonstrate that inclusion of externally-sequenced samples can pose analytical challenges. Specifically, we show that using ancestry-matched controls from publicly-available datasets can strongly inflate association test statistics if the sequences have not been processed together from BAM-level data. Sequence data can contain underlying structure due to sequencing platforms, coverage, alignment and calling algorithms. Inclusion of externally sequenced cases and controls (sequenced either separately or together) is non-trivial must be approached with extreme rigor. Methods for handling the merging and calling of heterogeneous sequencing datasets are currently being applied in large consortia^31–33^ and indicate that obtaining a high-quality dataset requires calling genotypes from petabytes of raw data in a uniform way across the full set of samples. Additional developments in handling heterogeneous sequence data include association testing on read-level data to allow for externally sequenced controls.^34^ Although these methods come with significant computational burden, similar approaches could be applied in Project MinE, in an effort to expand beyond the 22,500 samples sequenced initially by the project.

### The future of Project MinE

With the veritable explosion in methods and approaches being developed for wholegenome sequence data analysis, Project MinE will seek to leverage new and powerful methods for uncovering ALS risk variants. Current plans include more sophisticated functional annotation techniques, such as unsupervised-learning approaches to discriminate between functionally relevant and benign variation in the genome^35,36^; implementing a linear mixed model approach to burden testing^30^; a burden testing framework that will focus not only on genes but also on regulatory and other non-coding elements; and analyses that investigate both variation and methylation data. We have additionally established a pipeline to lift all our data to the newest genome build (hg38), which will likely yield better coverage of the genome in our samples. As the size of the data expands and additional collaborators join the effort, we will explore additional data structures that can facilitate transcontinental data analysis with minimal physical movement of data.

Finally, in addition to testing SNVs and indels, we will explore the role of more complex genetic variation such as structural variants and repeat expansions, which are known to play an important role in ALS susceptibility, yet have never been studied at genome-wide scale. With a global collaboration in place, a wealth of genetic variation being generated, and new methods for sequence data constantly in development, Project MinE will be the largest and most complete study of ALS genetics to date, poised to reveal novel risk loci, fine-map known disease genes, and shed light on the biological drivers of disease.

## Funding

*Research leading to these results has received funding from the European Community’s Health Seventh Framework Programme (FP7/2007-2013)*.

### The Netherlands

This study was supported by the ALS Foundation Netherlands.

### Belgium

*University Hospital Leuven (KU Leuven)*

*The Vesalius Research Center (VIB Leuven) (Belgium)*

### Ireland

*Trinity College Dublin*

### Spain

*Hospital Carlos III Madrid*

### Turkey

This study was supported by the Suna and Inan Kirac Foundation and Bogazici University.

### United Kingdom

SN is supported by the National Institute for Health Research (NIHR) Biomedical Research Centre at South London and Maudsley NHS Foundation Trust and King’s College London and the NIHR University College London Hospitals Biomedical Research Centre, and by awards establishing the Farr Institute of Health Informatics Research at UCLPartners, from the Medical Research Council, Arthritis Research UK, British Heart Foundation, Cancer Research UK, Chief Scientist Office, Economic and Social Research Council, Engineering and Physical Sciences Research Council, National Institute for Health Research, National Institute for Social Care and Health Research, and Wellcome Trust (grant MR/K006584/1).

## References

1 Hardiman O, van den Berg LH, Kiernan MC. Clinical diagnosis and management of amyotrophic lateral sclerosis. Nat Rev Neurol 2011; 7: 639–49.

2 Al-Chalabi A, Fang F, Hanby MF, et al. An estimate of amyotrophic lateral sclerosis heritability using twin data. J Neurol Neurosurg Psychiatry 2010; 81: 1324–6.

3 Byrne S, Walsh C, Lynch C, et al. Rate of familial amyotrophic lateral sclerosis: a systematic review and meta-analysis. J Neurol Neurosurg Psychiatry 2011; 82: 623–7.

4 Rosen DR, Siddique T, Patterson D, et al. Mutations in Cu/Zn superoxide dismutase gene are associated with familial amyotrophic lateral sclerosis. Nature 1993; 362: 59–62.

5 Sreedharan J, Blair IP, Tripathi VB, et al. TDP-43 mutations in familial and sporadic amyotrophic lateral sclerosis. Science 2008; 319: 1668–72.

6 Vance C, Rogelj B, Hortobágyi T, et al. Mutations in FUS, an RNA processing protein, cause familial amyotrophic lateral sclerosis type 6. Science 2009; 323: 1208–11.

7 DeJesus-Hernandez M, Mackenzie IR, Boeve BF, et al. Expanded GGGGCC hexanucleotide repeat in noncoding region of C9ORF72 causes chromosome 9p-linked FTD and ALS. Neuron 2011; 72: 245–56.

8 Renton AE, Majounie E, Waite A, et al. A hexanucleotide repeat expansion in C9ORF72 is the cause of chromosome 9p21-linked ALS-FTD. Neuron 2011; 72: 257–68.

9 Elden AC, Kim H-J, Hart MP, et al. Ataxin-2 intermediate-length polyglutamine expansions are associated with increased risk for ALS. Nature 2010; 466: 1069–75.

10 van Rheenen W, Shatunov A, Dekker AM, et al. Genome-wide association analyses identify new risk variants and the genetic architecture of amyotrophic lateral sclerosis. Nat Genet 2016; published online July 25. DOI:10.1038/ng.3622.

11 van Es MA, Veldink JH, Saris CGJ, et al. Genome-wide association study identifies 19p13.3 (UNC13A) and 9p21.2 as susceptibility loci for sporadic amyotrophic lateral sclerosis. Nat Genet 2009;41:1083–7.

12 Fogh I, Ratti A, Gellera C, et al. A genome-wide association meta-analysis identifies a novel locus at 17q11.2 associated with sporadic amyotrophic lateral sclerosis. Hum Mol Genet 2014; 23:2220–31.

13 Nelson MR, Wegmann D, Ehm MG, et al. An Abundance of Rare Functional Variants in 202 Drug Target Genes Sequenced in 14,002 People. Science 2012; 337: 100–4.

14 Genome of the Netherlands Consortium. Whole-genome sequence variation, population structure and demographic history of the Dutch population. Nat Genet 2014; 46: 818–25.

15 Lee S, Abecasis GR, Boehnke M, Lin X. Rare-variant association analysis: study designs and statistical tests. Am J Hum Genet 2014; 95: 5–23.

16 Millar AP, Baranova T, Behrmann G, et al. dCache, agile adoption of storage technology. J Phys Conf Ser 2012; 396: 032077.

17 Brooks BR, Miller RG, Swash M, Munsat TL, World Federation of Neurology Research Group on Motor Neuron Diseases. El Escorial revisited: revised criteria for the diagnosis of amyotrophic lateral sclerosis. In: Amyotrophic lateral sclerosis and other motor neuron disorders : official publication of the World Federation of Neurology, Research Group on Motor Neuron Diseases. 2000:293–9.

18 Huisman MHB, de Jong SW, van Doormaal PTC, et al. Population based epidemiology of amyotrophic lateral sclerosis using capture-recapture methodology. J Neurol Neurosurg Psychiatry 2011; 82: 1165–70.

19 Gordon PH, Miller RG, Moore DH. ALSFRS- R. Amyotroph Lateral Scler Other Motor Neuron Disord 2004; 5: 90–3.

20 Raczy C, Petrovski R, Saunders CT, et al. Isaac: ultra-fast whole-genome secondary analysis on Illumina sequencing platforms. Bioinformatics 2013; 29: 2041–3.

21 Yang J, Lee SH, Goddard ME, Visscher PM. GCTA: a tool for genome-wide complex trait analysis. Am J Hum Genet 2011; 88: 76–82.

22 Delaneau O, Marchini J, Zagury J-F. A linear complexity phasing method for thousands of genomes. Nat Methods 2012; 9: 179–81.

23 Browning BL, Browning SR. Improving the accuracy and efficiency of identity-by-descent detection in population data. Genetics 2013; 194: 459–71.

24 Purcell S, Neale B, Todd-Brown K, et al. PLINK: a tool set for whole-genome association and population-based linkage analyses. Am J Hum Genet 2007; 81: 559–75.

25 Chang CC, Chow CC, Tellier LC, Vattikuti S, Purcell SM, Lee JJ. Second-generation PLINK: rising to the challenge of larger and richer datasets. Gigascience 2015; 4: 7.

26 Wang K, Li M, Hakonarson H. ANNOVAR: functional annotation of genetic variants from high-throughput sequencing data. Nucleic Acids Res 2010; 38: e164.

27 Lin D-Y, Tang Z-Z. A general framework for detecting disease associations with rare variants in sequencing studies. Am J Hum Genet 2011; 89: 354–67.

28 Mathieson I, McVean G. Differential confounding of rare and common variants in spatially structured populations. Nat Genet 2012; 44: 243–6.

29 Gymrek M, McGuire AL, Golan D, Halperin E, Erlich Y. Identifying personal genomes by surname inference. Science 2013; 339: 321–4.

30 Listgarten J, Lippert C, Heckerman D. FaST-LMM-Select for addressing confounding from spatial structure and rare variants. Nat Genet 2013; 45: 470–1.

31 Lek M, Karczewski KJ, Minikel EV, et al. Analysis of protein-coding genetic variation in 60,706 humans. Nature 2016; 536: 285–91.

32 McCarthy S, Das S, Kretzschmar W, et al. A reference panel of 64,976 haplotypes for genotype imputation. Nat Genet 2016; 48: 1279–83.

33 Ganna A, Genovese G, Howrigan DP, et al. Ultra-rare disruptive and damaging mutations influence educational attainment in the general population. Nat Neurosci 2016; published online Oct 3. DOI:10.1038/nn.4404.

34 Hu Y-J, Liao P, Johnston HR, Allen AS, Satten GA. Testing Rare-Variant Association without Calling Genotypes Allows for Systematic Differences in Sequencing between Cases and Controls. PLoS Genet 2016; 12: e1006040.

35 Ionita-Laza I, McCallum K, Xu B, Buxbaum JD. A spectral approach integrating functional genomic annotations for coding and noncoding variants. Nat Genet 2016; 48: 214–20.

36 Ioannidis NM, Rothstein JH, Pejaver V, et al. REVEL: an Ensemble Method for Predicting the Pathogenicity of Rare Missense Variants. Am J Hum Genet 2016; published online Sept 21. DOI:10.1016/j.ajhg.2016.08.016.

37 Danecek P, Auton A, Abecasis G, et al. The variant call format and VCFtools. Bioinformatics 2011;27:2156–8.

